# Genomic analysis uncovers unique haplotype signatures from subspecies and agronomic types associated with blanchability in groundnut

**DOI:** 10.1101/2025.06.21.660844

**Authors:** Priya Shah, Sunil S. Gangurde, Ragavendran Abbai, Ramachandran Senthil, D. Khaja Mohinuddin, Madhvi Sharma, Prashant Singam, Kuldeep Singh, Pasupuleti Janila, Chuanzhi Zhao, Sandip K. Bera, Mei Yuan, Xingjun Wang, Rajeev K. Varshney, Manish K. Pandey

**Affiliations:** Center of Excellence in Genomics & Systems Biology (CEGSB) and Center for Pre-Breeding Research (CPBR), International Crops Research Institute for the Semi-Arid Tropics (ICRISAT), Hyderabad, Telangana-502324, India; Department of Genetics, Osmania University, Hyderabad, Telangana-500007, India; Institute of Crop Germplasm Resources, Shandong Academy of Agricultural Sciences (SAAS), Jinan-250100, China; ICAR, Indian Institute of Groundnut Research (IIGR), Junagadh, Gujarat-362001, India; Shandong Peanut Research Institute (SPRI), Chinese Academy of Agricultural Sciences (CAAS), Qingdao-266100, China; WA State Agricultural Biotechnology Centre, Centre for Crop and Food Innovation, Food Futures Institute, Murdoch University, Murdoch, Western Australia-6150, Australia

**Keywords:** blanchability, groundnut, haplotypes

## Abstract

Blanchability, defined as the ease of seed coat removal after roasting, is a vital trait for enhancing processing efficiency and product quality in groundnut (*Arachis hypogaea* L.). To enable a comprehensive haplotype-level genetic dissection of this trait, SNPs derived from whole-genome resequencing (WGRS) were used to perform genome-wide association study (GWAS) using multi-locus models (BLINK and FarmCPU) on a diverse groundnut mini-core collection phenotyped across two crop seasons. A total of 26 significant SNP-trait associations (STAs) were identified across multiple chromosomes, with major loci on chromosomes Ah05, Ah06, and Ah17, some of which were further validated using KASP (Kompetitive Allele-Specific PCR) markers. Candidate genes, such as those involved in cell wall biosynthesis (e.g., *galactoside 2-alpha-L-fucosyltransferase-like protein, protein kinase superfamily* members, and *glycerophosphoryl diester phosphodiesterase 3*), were found to be within linkage disequilibrium (LD) of the identified STAs, suggesting their plausible association with blanchability. Haplo-pheno analyses identified superior high-blanchability haplotypes; *Ah05HapBL3*, *Ah06HapBL5*, *Ah06HapBL10*, and *Ah17HapBL6, which* were predominantly found in the *fastigiata* subspecies (including Valencia and Spanish bunch agronomic types) from South Asia and South America, while the low-blanchability haplotypes were from *hypogaea* subspecies (including Virginia runner and Virginia bunch agronomic types) from Africa. Overall, this study provides valuable insights for customizing blanchability through haplotype-based breeding of processing-grade cultivars thereby improving groundnut value chains to meet diverse industrial demands.

## 1. Introduction

Groundnut (*Arachis hypogaea* L.; 2n = 4x = 40) is a globally important oilseed and economic crop of multi-purpose utility [1]. Groundnut genotypes can be broadly categorized into two subspecies: *hypogaea* and *fastigiata,* and six distinct botanical varieties: *hypogaea*, *hirsuta*, *fastigiata*, *vulgaris*, *aequatoriana*, and *peruviana* [2,3]. Additionally, groundnut accessions exhibit a remarkable array of morphological diversity, grouped into four agronomic types; Virginia runner, Virginia bunch, Valencia bunch, and Spanish bunch. Each agronomic type exhibits distinct growth habits, plant architecture, pod morphology, seed size, maturation duration, and yield performance. The Virginia runner and Virginia bunch types, belonging to subspecies *hypogaea*, are characterized by an absence of flowers on the main stem, an alternating pattern of vegetative and reproductive nodes on lateral branches, and a predominantly spreading or semi-erect growth habit. Virginia bunch types generally produce larger seeds and mature earlier than runners. In contrast, the Spanish and Valencia bunch types, under subspecies *fastigiata*, exhibit erect or compact growth habits, the presence of flowers on the main stem, sequential reproductive nodes, and smaller seed sizes. While the Spanish types are known for high oil content and early maturity, the Valencia types typically display three-seeded pods and are suitable for direct consumption, for example, such as nuts [4].

Groundnut products, such as salted, raw, or roasted nuts, oil, butter, candies, and groundnut flour, hold significant industrial importance. Here, ‘blanching’; the removal of seed coat is amongst the most critical steps to enhance the quality and shelf life of groundnut-based products [5]. As a result, ‘blanchability’, which refers to the ease with which the seed coat is removed, has become a key trait for groundnut industries in recent years. Blanching is a multistep process involving drying, roasting, mechanical rubbing between hard and soft surfaces, and air blowing to remove the seed coat. Several devices such as laboratory blanchers (American Society of Agricultural and Biological Engineers, 2006), have been developed to facilitate this process [5–7]. Improper removal of seed coats, and germ remnants impart a bitter taste to groundnut products, as, for example in groundnut butter, thus adversely impacting their industrial and economic value [5]. The genotypes and breeding lines possessing >60% blanchability are desirable for the commercial purpose such as groundnut butter, snack food, snack bars, groundnut flour and are considered processing-grade; however, commercial buyers often use 50% as a minimum threshold for acceptance [8]. The blanchability percentage requirement varies according to the manufacturers and product demand; for instance, genotypes with high split seeds are more suitable for candies, groundnut butter production, while low-blanchability is preferred for beer nuts and seed products with various confectionary coatings, where testa retention is desirable for added texture or adherence of coatings. Hence, both high and low-blanchability are valued in different industrial contexts, underscoring the importance of aligning breeding objectives with specific market requirements. This dual demand highlights the need for a nuanced approach in trait selection and cultivar development that considers the intended processing application.

Blanchability is influenced by a complex interplay of genetic, physiological, and environmental factors [9]. The groundnut seed coat, derived from the ovular integuments, is associated with multiple traits, including days to maturity, seed size, weight, and blanchability. Its properties are determined by both physical (thickness, density, fissures, cavitation) and chemical (lignin, polyphenols, alkaloids, anthocyanins) attributes [10,11]. Blanchability, in particular, is influenced by various factors such as genotype, kernel grade, post-harvest storage temperature, moisture content, storage conditions, and the thermal and hygroscopic properties of the cotyledon-skin interface [5,12,13]. The requirement of a large sample size (200 g) was a major issue for phenotyping blanchability; however, the previous findings, have suggested that a 50 g sample size should be sufficient for reliably estimating the blanching percentage [9,14].

In the past, bi-parental genetic mapping using the QTL-Seq approach (Middleton × Sutherland cultivar) reported genomic regions on chromosome A06 and B01 with R^2^ value of 10.3% and 5.6%, respectively [15]. Furthermore, studies using the U.S. mini-core (USMC) germplasm have investigated phenotypic variation and genotype × environment interactions to better understand and enhance the genetic improvement of blanchability [14]. Similarly, association mapping in the ICRISAT mini-core collection using SNP array has revealed genomic regions linked to blanchability; however, limited marker density and resolution have restricted the precision of these discoveries [9]. However, despite its critical industrial importance, a comprehensive blanchability-related genetic framework and haplotype signatures underscoring its selection pattern in diverse groundnut agronomic types remain largely elusive, thus hampering informed genetic improvement in breeding programmes.

Here, we utilized the ICRISAT groundnut mini-core collection [16] comprising 184 groundnut genotypes (57 Spanish bunch, 35 Valencia bunch, 48 Virginia bunch, and 33 Virginia runner). This collection was phenotyped for blanchability percentage across two crop seasons using the protocol by Janila et al., [14]. The availability of whole-genome resequencing (WGRS) datasets facilitated multi-locus genome-wide association study (GWAS) approaches using Bayesian-information and Linkage Disequilibrium Iteratively Nested Keyway (BLINK) and Fixed and Random Model Circulating Probability Unification (FarmCPU) to identify significant SNP-trait associations (STAs) for blanchability. Additionally, we developed diagnostic markers to accurately select for blanchability in a diverse groundnut mini-core collection developed by [16]. Furthermore, we identified key superior high-blanchability haplotypes underlying identified STAs, which provided detailed insights into their geographical distribution, as well as their current selection trends among various agronomic types. Taken together, the present study aims to provide a roadmap for groundnut haplotype-based breeding strategies using novel genetic donors to customize blanchability levels, thereby enhancing processing efficiency and meeting the diverse demands of global groundnut markets.

## 2. Material and Methods

### 2.1. Plant material and experimental field design

The diversity panel used in this study comprised of 184 accessions from the groundnut mini-core collection developed at ICRISAT, India [16]. The seed material of mini-core was procured from ICRISAT genebank and this panel represents the global genetic diversity of the groundnut germplasm. This mini-core collection of 184 accessions includes accessions from six botanical types: *hypogaea*, *hirsuta*, *fastigiata*, *peruviana*, *aequatoriana*, and *vulgaris*. Also, it represents variability for four agronomic types: Spanish bunch (57 genotypes), Valencia bunch (35 genotypes), Virginia bunch (48 genotypes), and Virginia runner (33 genotypes) **(Supplementary Table S1)**. Field trials were conducted using an alpha-lattice design across two consecutive seasons: the 2022 rainy season (R) and the 2022–2023 post-rainy season (PR) at ICRISAT, Patancheru (17°31′48.00″N, 78°16′12.00″E). The sowing of each genotype was conducted in a 4-meter row with spacing of 10 cm between plants and 60 cm between rows. The experimental field was characterized by red soil with a pH range of 6.0 to 6.3. Recommended standard agronomic practices for groundnut cultivation were implemented throughout both growing seasons to ensure optimal crop growth. Following harvest, phenotypic observations related to blanchability were recorded [9] using sound mature seeds with higher grade for each replication [6]**(Supplementary Table S1)**.

### 2.2 Phenotyping and data analysis

All the genotypes of the mini-core collection were phenotyped for blanchability using the protocol by Janila et al., [6]For this study, data was collected with two different sample sizes, 50 gm and 200 gm, with a blanching duration of 60 sec and 120 sec, respectively, at 30.00 rpm [9] using blancher, designed at ICRISAT. The weight of the blanched seeds and splits was recorded, and blanching percentage calculated using the formula below:

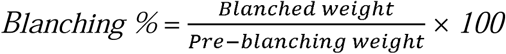

The phenotypic variation observed in the mini-core collection of blanchability was analysed with 50gm rainy season (50_R), 50gm post-rainy season (50_PR), 50gm_rainy_post-rainy (50R_PR), 200gm_post-raniy (200_PR) [9] **(Supplementary Table S1)** (**Figure 1**).

**Figure 1.**
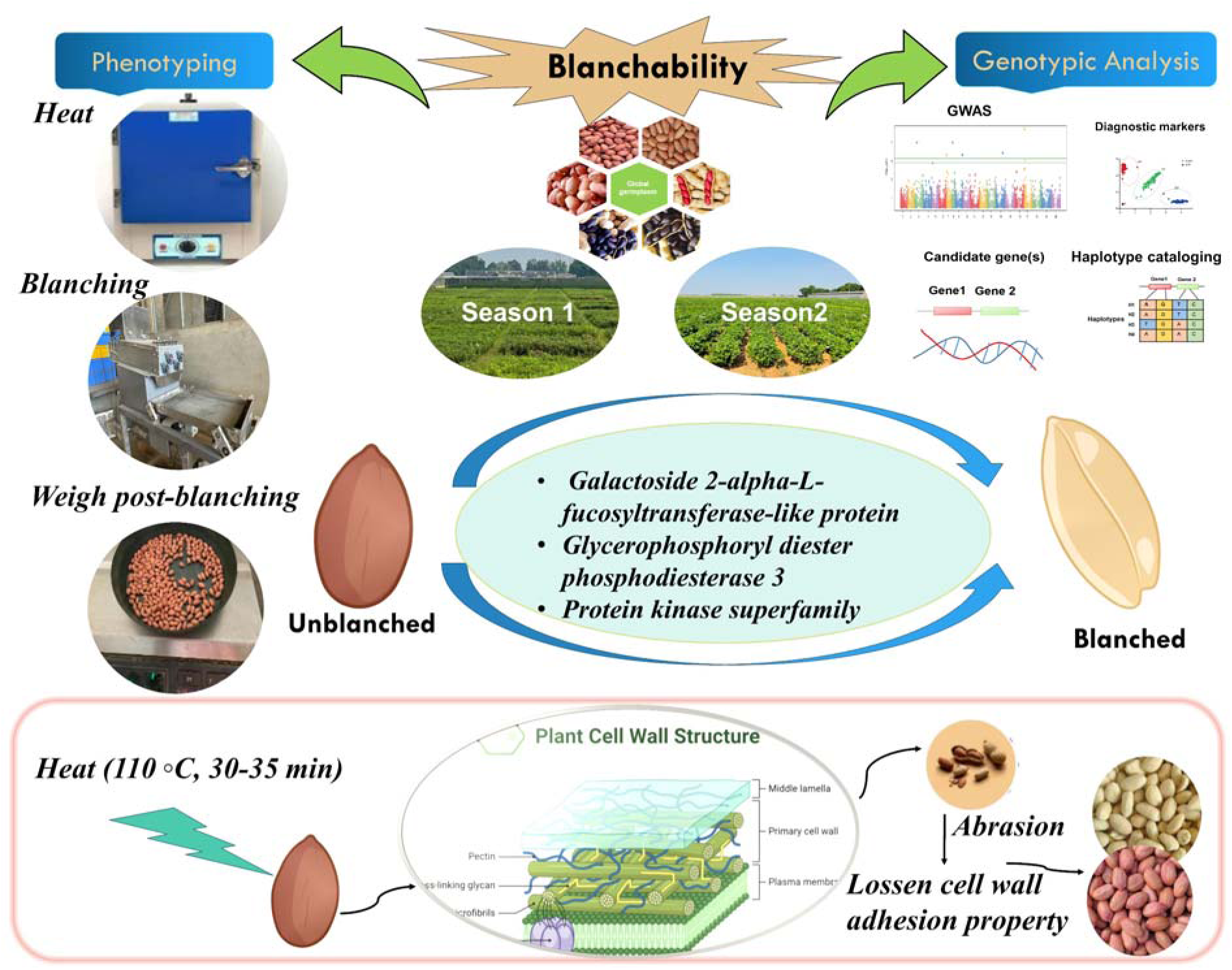
Phenotyping and genotyping for groundnut blanchability. Blanchability analysis in groundnut mini-core set to identify the marker and gene associated with the trait.

### 2.3 DNA extraction, sequencing and SNP calling from WGRS data

Genomic DNA for each genotype was extracted using Nucleospin Plant II kit (Macherey-Nagel, Düren, Germany). 100 mg tender leaf tissue was used for each genotype to extract high quality genomic DNA. Whole-genome resequencing was conducted using the Illumina sequencing platform, achieving an average depth of 10× coverage per accession, thereby enabling reliable variant detection and comprehensive genome-wide analysis. Post-sequencing, the adapter sequences were initially trimmed followed by filtering out the low-quality reads i.e., with > 20% low-quality bases (quality value ≤ 7) and > 5% “N” nucleotides using SOAP2 to ensure high-quality sequencing data for further analysis. The clean sequence reads were then mapped onto the reference genome of cultivated tetraploid “Tifrunner” v.2.0 [17] using SOAP2 software with parameters “-m 300 -x 600 -s 35 -l 32 -v 5 -p 4” followed by calculation of the likelihood of all possible genotypes for each sample using SOAPsnp3 to get the maximum likelihood estimation of the allele frequency in the population [17]. The low-quality variants were then filtered-out using the stringent criteria for sequencing depth (>10000 and <400), mapping times (>1.5), and quality score (<20); and at the end, the SNP loci with estimated allele frequency ≠0 or 1. In addition, SNPs with ≥50% missing among genotypes were then further filtered-out and remaining high quality SNPs were considered for further analysis (**Supplementary Figure S1).**

### 2.4 Principal component analysis and GWAS for identifying SNPs-associated with blanchability

Population structure was estimated using principal component analysis (PCA) based on a genetic distance matrix (Roger’s matrix) to identify genetic clusters with similar genetic backgrounds within mini-core collection. PCA analysis was conducted using the SelectionTools package in R [18] **(Supplementary Figure S2).** Linkage disequilibrium (LD) decay in the groundnut mini-core diversity panel was analyzed using PopLDdecay v.3.29 [19] and the SelectionTools package in RStudio [18] **(Supplementary Figure S3 (A) and (B)).** To address population structure and relatedness in the GWAS, both the Q matrix (from population structure analysis) and the K matrix (pairwise kinship estimates) were incorporated into the multi-locus model. Together, these matrices effectively manage confounding effects, reducing false-positive associations that could arise from shared ancestry or familial relationships. Quantile-Quantile (Q-Q) plots were well visualized by plotting observed -log10 (*p*) values against expected -log10 (*p*) values for genome-wide SNPs, using the CMplot package in RStudio. Manhattan plots were also visualized using CMplot [20]. To correct spurious associations in the GWAS, a Bonferroni correction was applied at a 5% significance level. The threshold for significant marker-trait associations was determined using the formula α*/n*, where α is the overall significance level, and *n* represents the total number of SNPs used in the GWAS analysis. For instance, the Bonferroni threshold (*p-value)* = 4.97 × 10. The percentage of phenotypic variance explained (PVE) by significant SNPs was obtained from the models used, with PV for each SNP calculated as the squared correlation between phenotype and genotype. The significance threshold for STAs in WGRS-based GWAS analysis was set at [–log10 (*p*) >6], ensuring the selection of reliable SNPs (**Figure 2; Supplementary Figure S4) (Table 1)**. To identify STAs, GWAS analysis was performed using the multilocus model, FarmCPU and BLINK implemented in Genome Association and Prediction Integrated Tool (GAPIT) package using the RStudio [21]. Multi-locus models BLINK and FarmCPU incorporate both population structure (Q) and kinship (K) matrices during association mapping. Further, the effects of alleles underlying significant and stable SNP markers were analyzed using methods described by Su et al., and Alemu et al., [22,23]. Genotypes were grouped based on specific SNP alleles, and mean differences were evaluated using Tukey’s HSD test **(Supplementary Figure S6)**. Further, the identified STAs were validated, for this, the DNA was extracted from the l00 mg leaf tissue of 25 samples (10 genotypes and 15 breeding lines) and were included in the marker validation panel.

**Figure 2:**
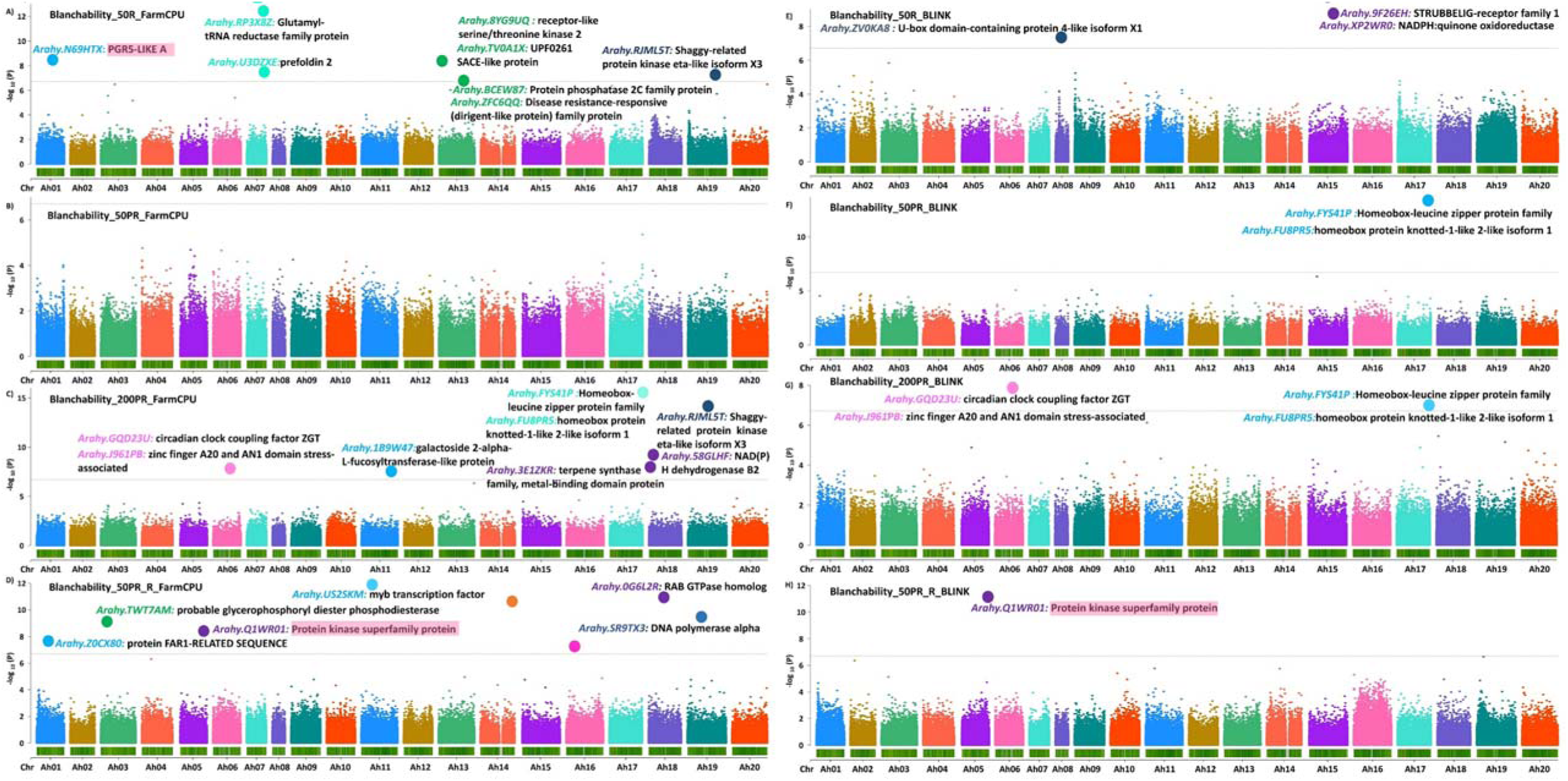
Manhattan plots representing the identified STAs associated with the blanchability on the basis FarmCPU (A, B, C, D) and BLINK (E, F, G, H) model in GAPIT: A,E: Blanchability 50R (50gm sample size in rainy season), B,F: Blanchability 50PR (50gm sample size in post-rainy season), C,G: Blanchability 200PR (200gm sample size in post-rainy season) and D,H: Blanchability 50PR_R (50gm sample size in post-rainy and rainy season). In these Manhattan plots, the highlighted (pink) region shows the gene associated with the validated polymorphic markers on chromosome Ah01 and Ah05; respectively. The other dots represent the significant STAs and the genes found to be associated with blanchability.

**Table 1:**
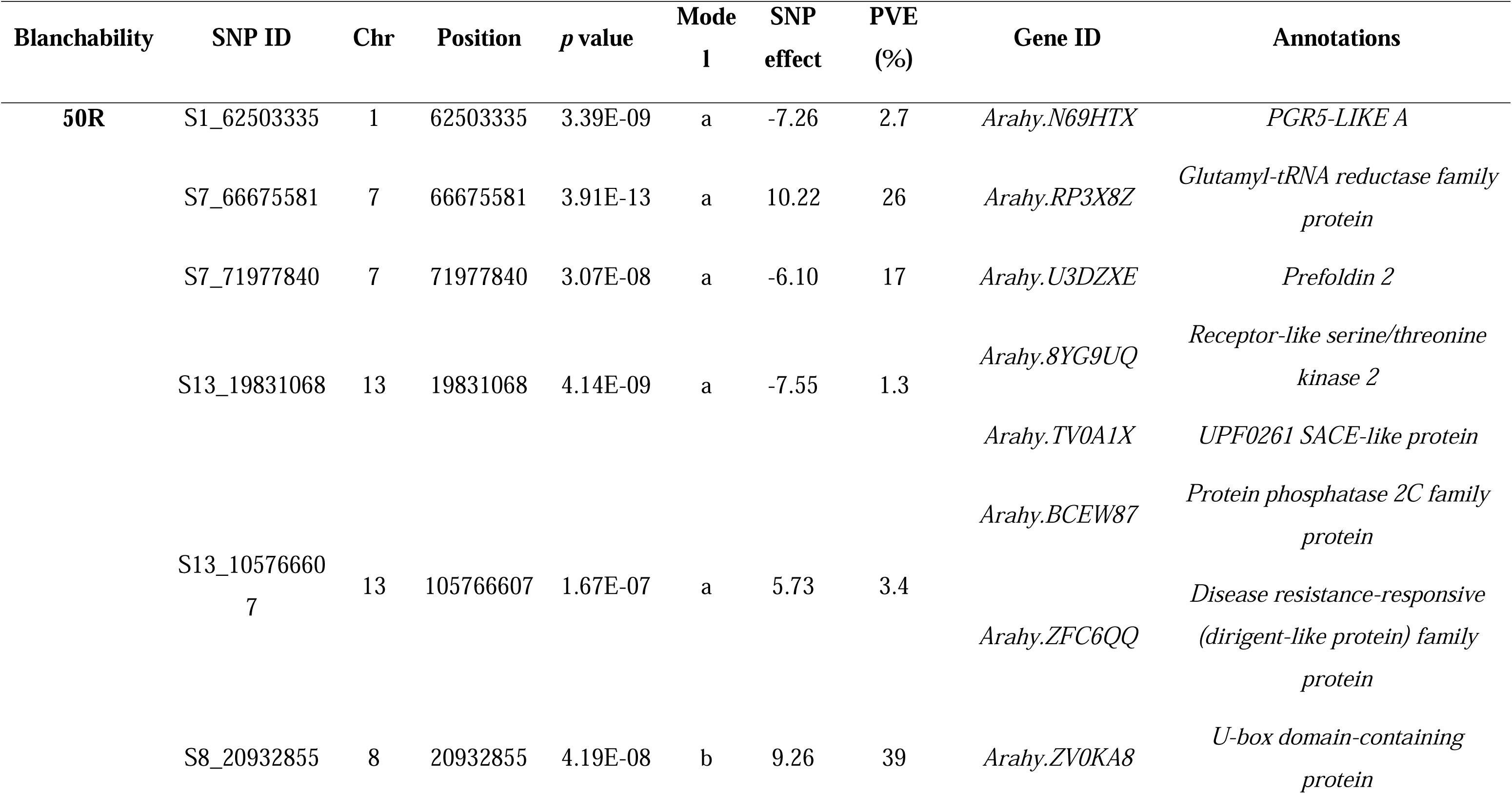
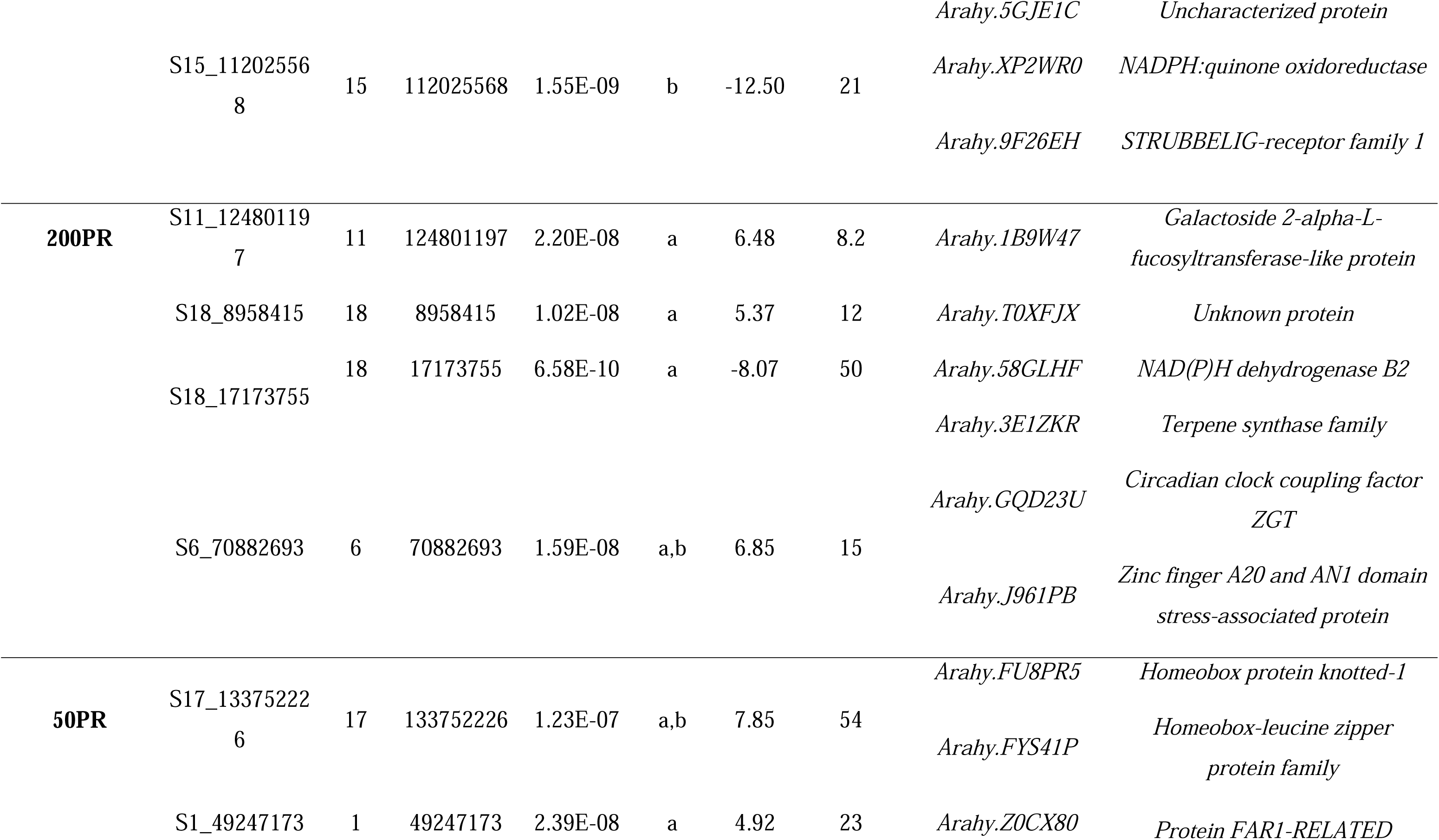
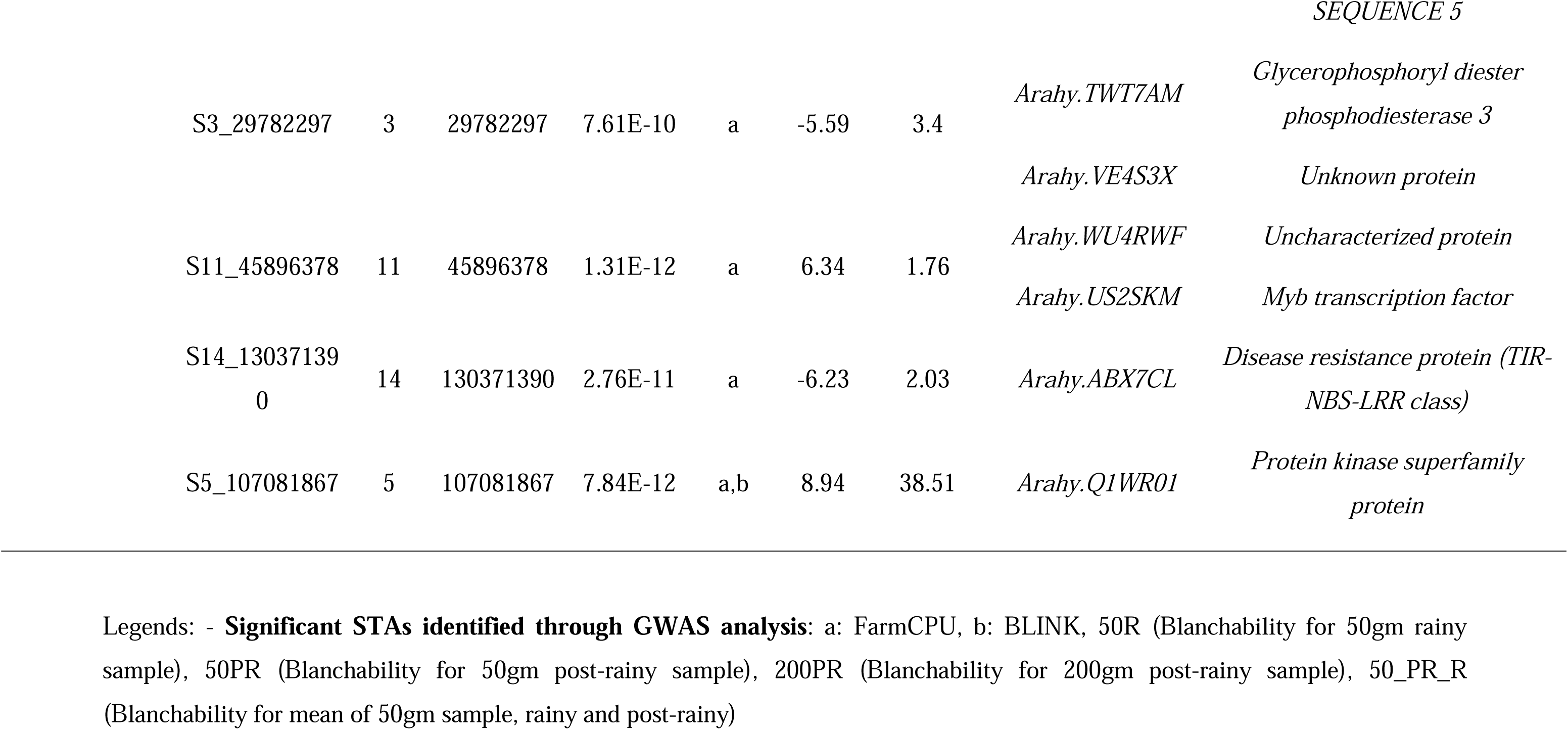
Significant SNP trait associations and corresponding candidate genes identified for groundnut blanchability.

### 2.5 Haplo-pheno analysis and identification of superior haplotypes for blanchability

For each significant STA identified from the GWAS study, a subset of up to ten representative SNPs that are in LD with the lead SNP was first selected. These representative SNPs were chosen to span a spectrum of LD levels with the significant SNP: i. low LD (r^2^ > 0.30 and <= 0.50); ii. SNPs with moderate LD (r^2^ > 0.50 and <= 0.70), and iii. high LD (r^2^ > 0.70). This LD binning approach identified the SNPs with varying degrees of correlation, capturing distinct haplotype block patterns around the lead SNP. The genotypic combinations of these representative SNPs were then used to define haplotype groups, such that all available unique patterns within the panel were comprehensively represented. FlapJack software was used to visualize the identified haplotype groups [24]. Subsequently, to assess the phenotypic impact of each haplotype, ‘haplo-pheno’ analysis was performed. This involved multiple range comparisons of the phenotypic means among all the haplotype groups, which comprised at least three genotypes. One-way ANOVA, followed by Tukey’s HSD post-hoc test, was used to determine statistical significance. The haplo-group(s) with the most optimal/preferred phenotypic mean values were designated as superior haplotype(s).

### 2.6. Development of KASP-based markers

GWAS analysis identified 26 significant STAs, of which 6 highly significant markers were selected for the development of KASP markers **(Supplementary Figure S5).** The selected polymorphic SNPs were in genomic regions proximal to candidate genes distributed across three different chromosomes. Further, KASP markers were designed based on 50-base pair upstream and downstream flanking sequences. For each SNP, two allele-specific forward primers and one common reverse primer were synthesized at Intertek Pvt. Ltd., Hyderabad, India. The designed KASP markers were validated using a panel of 25 samples, comprising five high and five low-blanchability genotypes as well as 15 breeding lines with varying blanchability from ICRISAT (**Figure 4**).

### 2.7 Identification of candidate genes for superior blanchability

Peanut base webtool (https://Peanutbase.org/) was used to identify the candidate genes using GBrowse (cultivated peanut) version 1. The candidate genes corresponding with the STAs were identified using peanut base reference genome sequence annotations (www.peanutbase.org). The physical positions of the SNP were mapped onto the *Arachis hypogaea* reference genome v1.0 available in PeanutBase (www.peanutbase.org) and the corresponding genes were identified. The SNP subsiding start and ending position of the gene or exons were explored for the candidate gene discovery based on their biological function annotation related to the trait of interest **(Table 1**). Candidate genes involved in regulating cell wall integrity and adhesion, previously reported in the literature, were described in the context of blanchability.

## 3. Results

### 3.1 Phenotyping variability for blanchability in ICRISAT mini-core set

We have generated and used data from two seasons in two replications with different sample size; 50gm rainy season (50_R), 50gm post-rainy season (50_PR), 50gm_rainy_post-rainy (50R_PR), 200gm_post-raniy (200_50PR), as phenotypes for GWAS analysis. The highest blanchability noted in the high-blanchability lines was ranging upto 80% while in low the blanchability was observed to be 3.98 as the lowest [9] **(Supplementary Table S1)**.

### 3.2. Characterization of variants, principal component analysis and linkage disequilibrium decay

A total of 5,61,099 SNPs were identified using WGRS approach. Minor allele frequency (MAF) was applied at 0.05 level. A working subset of 2,55,144 filtered SNPs with 0.2% heterozygosity, and 0.05% MAF was used for GWAS analysis with bonferroni correction of 5%, across 167 accessions were subsequently employed for association mapping studies. SNP (Single Nucleotide Polymorphism) density across chromosomes, segmented into 1Mb windows, shows the overall SNP counts (**Supplementary Figure S1)**.

Heatmaps and dendrograms of the kinship matrix, based 2,55,144 on polymorphic SNPs for the studied genotypes, for WGRS data, indicated that there was clear clustering of genotypes in the mini-core collection **(Supplementary Figure S2; S3 (A) and (B))** (**Supplementary Table S2).** From WGRS data, the population structure based on groundnut sub-species, botanical variety and agronomic type also revealed two distinct clusters: cluster1 (88 genotypes): *fastigiata*, Valencia bunch/ *hypogaea*, Virginia runner/ *vulgaris*, Spanish bunch while for cluster2 (72 genotypes): *hypogaea*, Virginia bunch or Virginia runner/ *hirsuta*/ *aequatoriana*/ *peruviana*/ *vulgaris*, Spanish bunch/ *fastigata*, Valencia bunch **(Supplementary Figure S3 (A) and (B))** (**Supplementary Table S2).** The graphical representation of the linkage disequilibrium characteristics of the mini-core collection is presented in **Supplementary Figure S3 (A) and (B);** respectively. For WGRS data, the average r^2^ value of the genome was 0.2, and the LD decay was found to initiate at an r^2^ value of 0.70, and reached half-decay at 0.28. The LD decay curve intersected with the half decay at 50 Kb, which represents the genome-wide critical distance to detect linkage. Hence, candidate genes associated with markers identified for blanchability within this distance were considered as associated candidate genes.

### 3.3. Genomic regions associated with blanchability in groundnut

GWAS was performed using 2,55,144 high-quality SNPs with less than 20% missing data. These SNPs were distributed across the twenty chromosomes of groundnut. SNP genotyping data of 2,55,144 SNPs along with information on population structure and kinship matrix were used for genome-wide association analysis for blanchability with the phenotypic data obtained from the 2022 rainy season and 2022-2023 post-rainy season. After the calculation, the Bonferroni correction threshold level of -log (10) ‘*p*’ value was set to 6.0 for analysis with WGRS data. Deviations in *p*-values at the initial stage indicated the presence of population stratification. The FarmCPU and BLINK models were found to significantly affect the detection of associations contributing to the traits of interest **(Figure 2; Supplementary Figure S4).**

A total of 26 STAs were found with the two different models used; 8 STAs were associated with 200PR, in which the range of ‘*p*’ value; 2.53 × 10^-16^ to 1.78 × 10^-7^ and PVE; 0.86 to 54.03%. While for data set with 50PR_R mean (9 STAs), 50PR (1 STA), 50R (8 STAs), the ‘*p*’ values ranged from 1.31× 10^-12^ to 5.77 × 10^-8^ with PVE range from 0 to 38.51%, the ‘*p*’ values ranged from 1.34 × 10^-8^ with PVE value 43.12%, the ‘*p*’ values range from 3.91 × 10^-13^ to 1.67 × 10^-7^ with PVE range from 0 to 38.75%. Among the STAs identified, the STA on chromosome Ah17 exhibited the maximum PVE value of 54.03% **(Table 1)**. The 6 polymorphic markers that were identified for blanchability from the analysis with the 4 different data-sets, were distributed across chromosome Ah01 (1), Ah05 (1), Ah06 (1), Ah07 (1), Ah16 (1), and Ah17 (1) **(Supplementary Figure S5).** This include *S1_62503335* (Ah01) (50_R), *S5_107081867* (Ah05) (50_PR_R), *S6_70882693* (Ah06) (200_PR), *S7_66675581* (Ah07) (50_R), *S16_34347730* (Ah16) (50_PR_R) and *S17_133752226* (Ah17) (200PR, 50PR). Further, we have determined the allelic effects and allelic distribution of these stable STAs (Supplementary Figure S6).

### 3.4. Candidate genes associated with blanchability

The total number of genes in the groundnut genome has been estimated to be around 83,709. This count is based on the sequenced groundnut genome and the identified gene models within it [17]. The SNPs obtained from the analysis were compared against the groundnut genome sequence using physical positions. The functions of the gene underlying the respective SNP was investigated using Peanut base. It was observed that 15 SNPs were found to possess candidate genes that were associated with seedcoat and cell wall adhesion property, that influence blanchability **(Table 1)**. This includes genes*, galactoside 2-alpha-L-fucosyltransferase-like protein (Arahy.1B9W47), glycerophosphoryl diester phosphodiesterase3-like (Arahy.TWT7AM), Protein kinase superfamily protein (Arahy.Q1WR01).* The gene *Arahy.1B9W47* was identified from *S11_124801197* (Ah11), explaining 8.2% of phenotypic variation (PVE), while *Arahy.TWT7AM* was associated with marker *S3_29782297* (Ah03), accounting for 3.4% PVE. The *Arahy.Q1WR01* gene, linked to *S5_107081867* (Ah05), explained the PVE at 38.51% and possess a validated SNP (snpAH00569), highlighting its potential significance in controlling blanchability traits.

### 3.5. Haplo-pheno analysis of STAs for the identification of superior haplotypes for blanchability

The availability of WGRS dataset permitted a comprehensive genomic investigation at haplotype resolution. Notably, to determine the impact of haplotype diversity on phenotypic performance, haplo-pheno analysis was conducted, where differences in the phenotypic means were correlated with the distinct haplotype signatures of all 26 blanchability-related STAs. Significant haplotype diversity *Ah05HapBL*, *Ah06HapBL*, and *Ah17HapBL* across 3 STAs; *S5_107081867* (Ah05) (50_PR_R), *S6_70882693* (Ah06) (200_PR), and *S17_133752226* (Ah17) (200PR, 50PR); were observed; respectively **(Figure 3, Supplementary Table S3-S11).** The objective was to identify superior haplotypes, defined as those leading to a significantly high percentage for blanchability compared to other haplotypes. Statistically validated by one-way ANOVA followed by Tukey’s HSD post-hoc test, all candidate loci possessing more than two haplotypes were scrutinized to pinpoint superior haplotypes **(Figure 3**).

**Figure 3.**
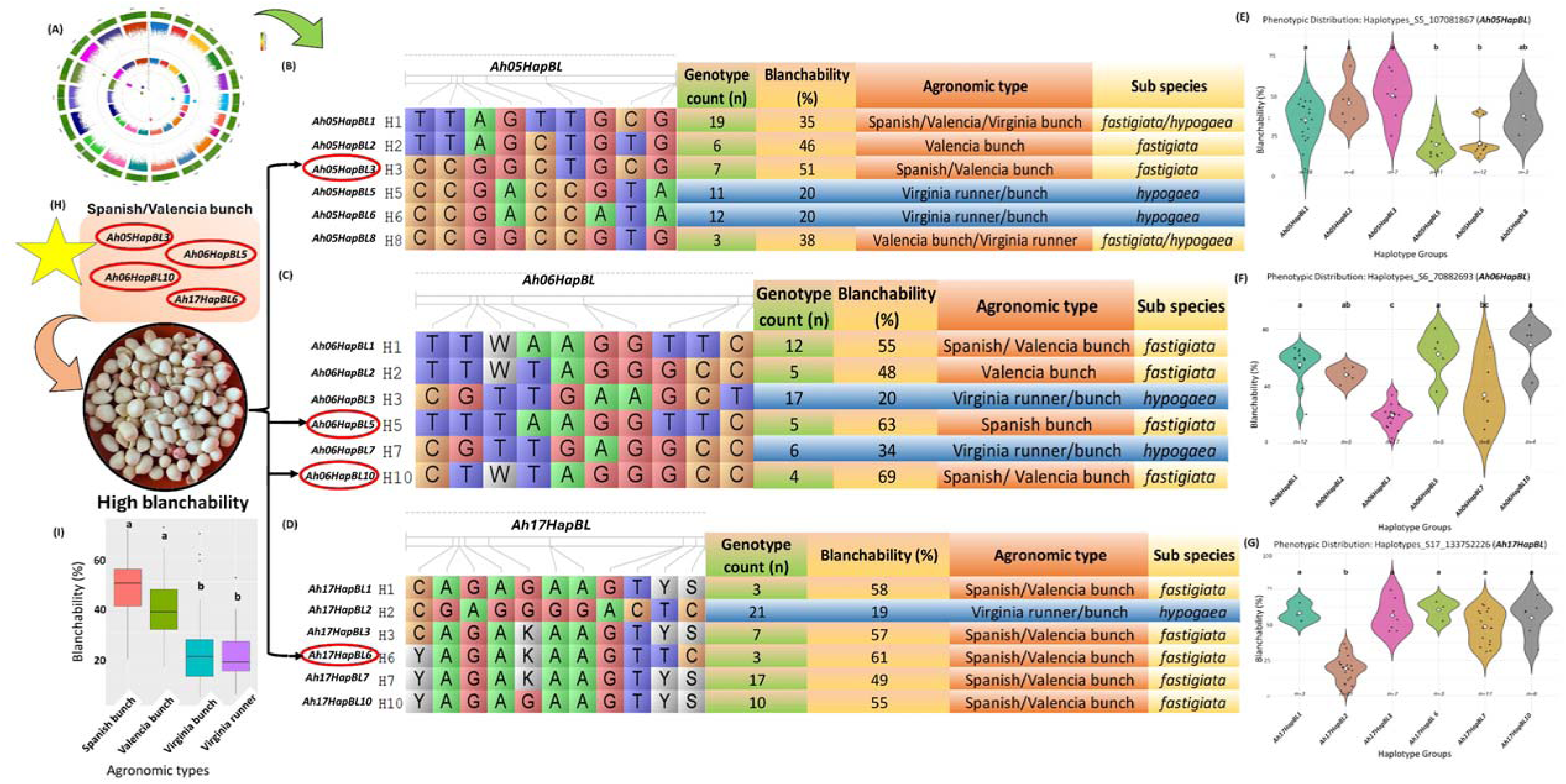
Superior haplotypes to tailor high blanchability in groundnut. (A) Significant STAs identified based on GWAS for blanchability. (B, C, D) LD-based haplotype (*Ah05HapBL*), (*Ah06HapBL*), and (*Ah17HapBL*) clustering of the genomic region (Ah05_107081867), (Ah06_70882693) and (Ah17_133752226); respectively, associated with blanchability. There were 24 haplotype groups in total of which 6 are depicted as they have at least 3 genotypes in their haplotype group. FlapJack software was used to visualize the haplotype groups. (E, F, G) Haplo-pheno analysis revealed that *Ah05HapBL3, Ah06HapBL5, Ah06HapBL10* and *Ah17HapBL6* confers high blanchability (>50%), while *Ah05HapBL5, Ah05HapBL6, Ah06HapBL3, Ah06HapBL7,* and *Ah17HapBL2* for low-blanchability (<35%). (H) The superior high blanchability haplotypes *Ah05HapBL3, Ah06HapBL5, Ah06HapBL10 and Ah17HapBL6* identified here can be used to get higher blanchability in groundnut. (I) High blanchability trait predominantly belongs to Spanish bunch and Valencia bunch agronomic type, while Virginia runner and bunch has low blanchability.

**Figure 4:**
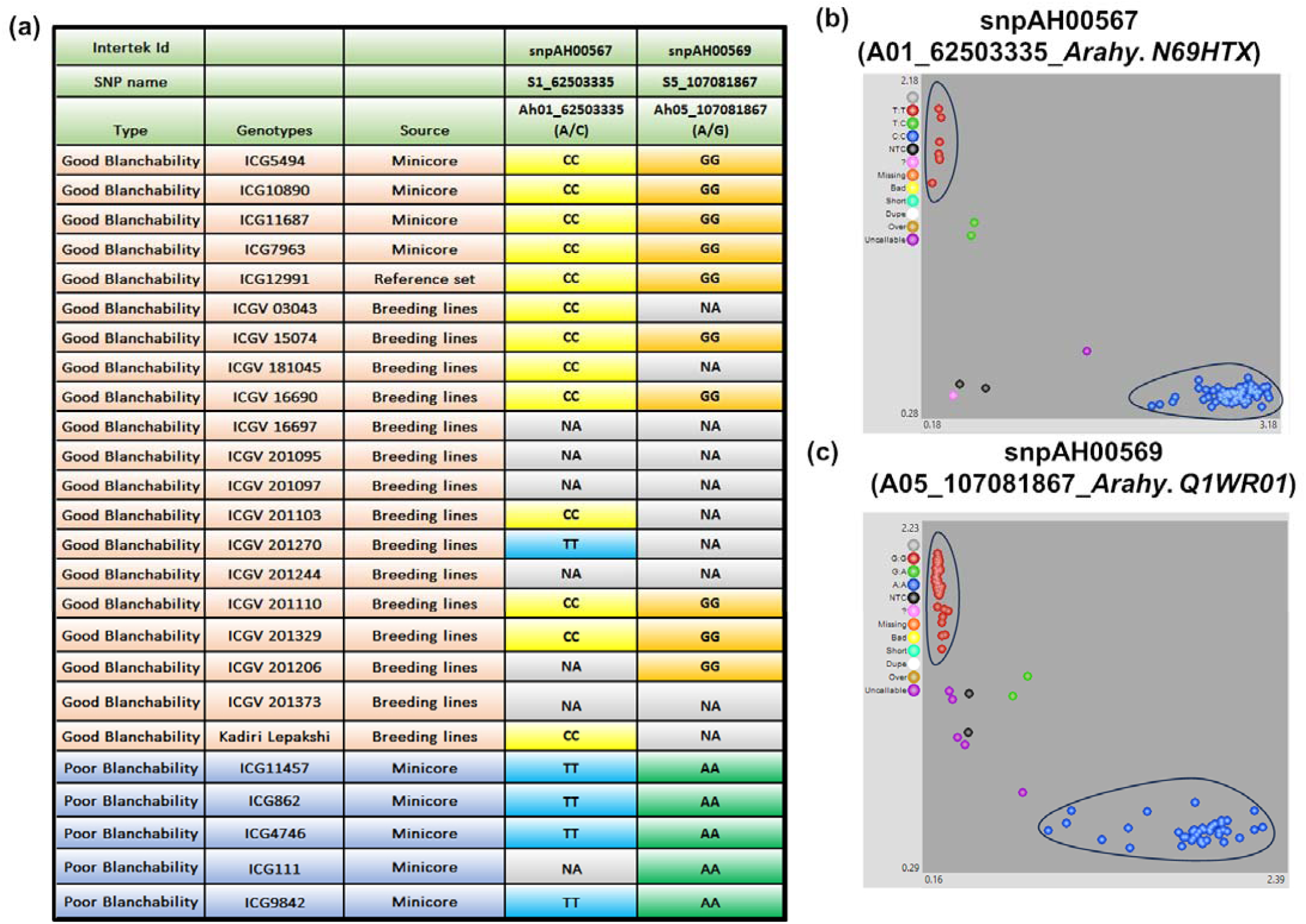
Development and validation of KASP markers from potential candidate genes identified for blanchability. (a) Validation panel includes 20 high-blanchability genotypes and groundnut varieties (50-70%-blanchability), and five low-blanchability genotypes (1-10%-blanchability). Two single-nucleotide polymorphisms (SNPs) show clear homozygous clusters for (b) snpAH00567 (Ah01_62503335) for *PGR5-like* protein (*Arahy.N69HTX*) gene, (c) snpAH00569 (Ah05_107081867) for *Protein kinase superfamily* (*Arahy.Q1WR01*) gene.

Four superior high-blanchability haplotypes (*Ah05HapBL3* (51%), *Ah06HapBL5* (63%)/ *Ah06HapBL10* (69%), *Ah17HapBL6* (55%)) were defined with blanchability >50%, while low-blanchability haplotypes are *Ah05HapBL5/ Ah05HapBL6*, *Ah06HapBL3 and Ah17HapBL2* with <20% blanchability, the high and low blanchability haplotypes belongs to haplotype group *Ah05HapBL, Ah06HapBL*, *Ah17HapBL;* respectively. The identified superior high-blanchability haplotypes, exhibited the highest mean blanchability in the respective group **(Figure 3**).

### 3.6. Development and validation of KASP markers for blanchability

To develop and validate diagnostic markers for blanchability, six SNPs located on Ah01, Ah05, Ah06, Ah07, Ah16 and Ah17 with one SNPs on each chromosome, and these SNPs were targeted for development of KASP markers. Primers were successfully developed for 6 SNPs; *S1_62503335* (TT/CC), *S5_107081867* (AA/GG), *S6_70882693* (AA/GG), *S7_66675581* (AA/GG), *S16_34347730* (CC/TT) and *S17_133752226* (AA/GG), and validated on a panel with contrasting blanchability **(Supplementary Figure S5).** The validation panel comprised of low and high-blanchability genotypes and breeding lines ranging from 6% to 80%. Of the six KASP markers selected for validation, two KASPs (*snpAH00567*, and *snpAH00569*) showed the expected polymorphism in the designed validation panel. Interestingly, these two polymorphic KASP markers differentiated genotypes and breeding lines with high and low-blanchability which belonged to genomic region detected on chromosome Ah01 and Ah05 **(Figure 4)**. Additionally, markers from chromosome Ah17 were also validated in few genotypes but majorly the calls were missing hence lacked confidence. Nevertheless, we have identified superior haplotypes which can be useful in breeding high-blanchability groundnut. The validated KASP markers accurately distinguished high and low-blanchability genotypes and breeding lines, and can be used as potential diagnostic markers for selecting blanchability breeding material at the very early stages of the varietal development process.

## 4. Discussion

Blanchability, defined as the ease of seed coat removal post-roasting, has emerged as a critical quality trait in breeding programs targeting edible and industrial applications [5]. The importance of this trait lies in its direct impact on processing efficiency and product quality, making it a key target in breeding programs. With the advancement in next-generation sequencing (NGS) technology, there has been increasing availability of various groundnut genomic resources, enabling the precise dissection of complex traits, including blanchability [25–27]. In this study, a mini-core diverse collection was analysed using WGRS and multi-environment phenotyping for blanchability. Our aim was to identify genomic regions, candidate genes and haplotype diversity for blanchability. We have utilized advanced models like FarmCPU, and BLINK in GWAS that significantly enhance reliability and precision, detecting a total of 26 significant STAs for blanchability **(Figure 2; Supplementary Figure S4, Table 1)**.

Haplo-pheno analysis of the identified STAs revealed significant haplotype diversity for blanchability, particularly at key loci on chromosomes Ah05 (*Ah05HapBL*), Ah06 (*Ah06HapBL*), and Ah17 (*Ah17HapBL*). This approach aims at a deeper understanding of the genetic factors influencing these economically important characteristics in groundnut cultivation. The *Ah17HapBL* was found to be associated with *S17_133752226* (Ah17), and exhibited the highest PVE (54.03%). The high-blanchability haplotypes associated with *Ah17HapBL* (*Ah17HapBL1* (mean blanchability 58%)*, Ah17HapBL3* (mean blanchability 57%)*, Ah17HapBL6* (mean blanchability 61%)*, Ah17HapBL10* (mean blanchability 55%)) predominantly originated from South Asia and South America, majorly belong to the sub-species *fastigiata*, botanical variety, *fastigiata/vulgaris*, and agronomic type, Valencia bunch/Spanish bunch. This underscores the influence of both genetic background and ecogeographic adaptation on processing quality. Similar association has been observed for the haplotypes associated with STAs *S5_107081867* (Ah05) and, *S6_70882693* (Ah06); where superior high-blanchability haplotypes, *Ah05HapBL3* (mean blanchability 51%) and *Ah06HapBL5* (mean blanchability 63%), *Ah06HapBL10* (mean blanchability 69%), belong to sub-species *fastigiata,* botanical variety, *fastigiata/vulgaris*, and agronomic type, Valencia bunch and Spanish bunch. Conversely, the clustering of low-blanchability accessions in *Ah05HapBL6* (mean blanchability 20%), *Ah06HapBL3* (mean blanchability 20%)*, Ah17HapBL2* (mean blanchability 19%) haplotype, predominantly from Virginia bunch and Virginia runner (sub-species *hypogaea*) types, particularly from Africa, highlights the challenges posed by genetic constraints within the *hypogaea* subspecies **(Figure 3**). These findings align with our previous study [9], that has revealed marked differences among the cultivated agronomic types of groundnuts, Spanish bunch genotypes consistently exhibited high-blanchability, whereas most Virginia runner types had low-blanchability. This indicates that the sub-species and agronomic type significantly impact blanchability, suggesting the influence of population structure on blanchability and more interestingly, a plausible genetic interaction with the loci regulating agronomic type, which remains to be elucidated. In this context, the superior high-blanchability haplotypes identified here can potentially be used to precisely customize blanchability in any genetic background **(Figure 5)**.

**Figure 5:**
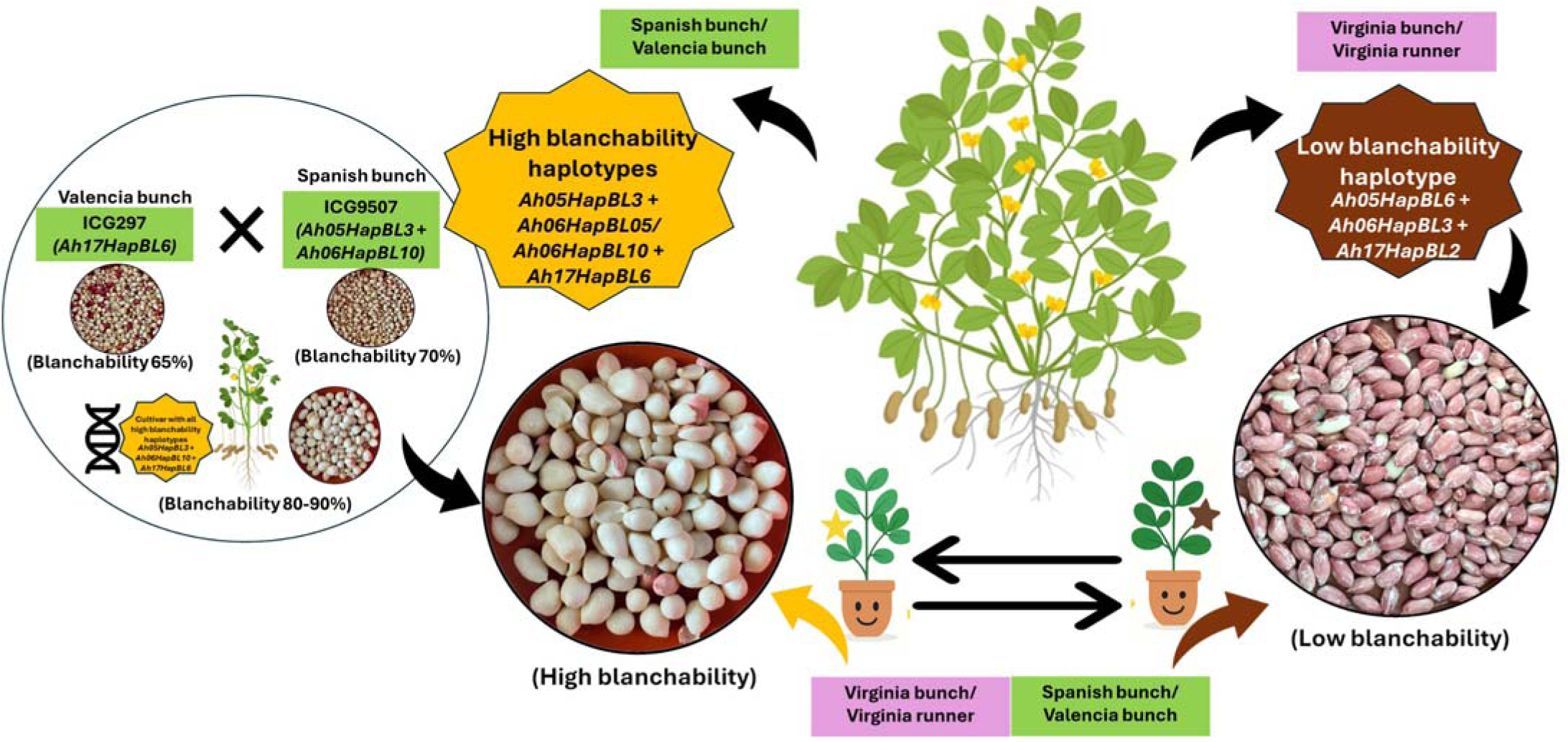
Haplotype-based breeding to customize blanchability into diverse groundnut cultivars. High-blanchability-associated haplotypes (*Ah05HapBL3, Ah06HapBL05/BL10, Ah17HapBL6*) (yellow star), are predominantly present in Spanish bunch and Valencia bunch groundnut types, whereas Virginia runner and Virginia bunch types commonly possess low-blanchability haplotypes (*Ah05HapBL6*, *Ah06HapBL3, Ah17HapBL2*) (brown star). The cross between high-blanchability genotypes (ICG9507 and ICG297) is expected to result in improved blanchability (80–90%). Overall, the identified superior high-blanchability haplotypes can be used to tailor high-blanchability in any genetic background, independent of the agronomic type, to match groundnut genotypes to the required industrial demand. This will enable the haplotype introgression from high-blanchability genetic backgrounds into low-blanchability types, enabling the development of Virginia-type cultivars with high-blanchability, and vice-versa.

Groundnut blanchability is a trait influenced by genetic and environmental factors. Previous QTL-seq in a recombinant inbred line (RIL) population derived from ‘Middleton’ × ‘Sutherland’ uncovered blanchability QTLs on chromosomes A06 (*Arahy.06_108,665,514*, *Arahy.06_108,812,907*) and B01 (*Arahy.11_15,264,657 Arahy.11_16,329,544*, *Arahy.11_18,994,278*), each explaining roughly 5.6–10.3 % of PVE and collectively accounting for 14.8 % of the trait variance [15]. Similarly, unpublished SNP-array work validated KASP markers for blanchability on chromosomes A01, A06, B04, and B07 [9]. In this WGRS based analysis, two KASP markers, *SnpAH00567* (*Ah01_62503335_Arahy.N69HTX*) and *SnpAH00569* (*Ah05_107081867_Arahy.Q1WR01*) were successfully validated, corroborating previous findings and emphasizing the repeatability of these loci across platforms [9].

Previous analysis with the 58K Axiom_*Arachis* SNP array detected a major STA on chromosome B07 that explained 35.3 % of the PVE [9]. In the current study, leveraging the higher marker density of WGRS, we have again identified a major significant STA on chromosome Ah17 with a PVE of 54.0 %. Moreover, the significant haplotype diversity was also identified for this STA and has revealed the superior high-blanchability haplotype for blanchability belonging to sub-species *fastigiata*, botanical variety; *fastigiata*/*vulgaris*, and agronomic type; Valencia bunch and Spanish bunch. However, for the STA associated with Ah17, in the KASP validation result showed it to be uncallable for maximum genotypes and breeding lines, which laid emphasis on gel-based marker for the validation of this particular STA. These findings indicate the significance of chromosome Ah17 to possess the key genomic region governing the blanchability. Additionally, the STAs on Ah05 (*S05_107081867)* also possess the significant haplotype diversity and has been validated successfully in the KASP validation as well.

Blanchability is linked to the seed coat characteristics, particularly adhesion and seed coat integrity, that determine how tightly the seed coat adheres to the cotyledons. In this study, most of the genes identified in association with significant STAs are directly tied to cell wall biosynthesis, modifications, remodeling and adhesion proteins Such *galactoside 2-alpha-L-fucosyltransferase-like protein*. In *Arabidopsis thaliana*, *galactoside 2-alpha-L-fucosyltransferase-like protein* specifically fucosylates xyloglucan, influencing cell wall integrity and potentially affecting cell adhesion [28]. Additionally, *glycerophosphoryl diester phosphodiesterase* has been identified to be involved crucially in cell wall organization [29].

However, for the genes *PGR5* and the *protein kinase superfamily*, which have been associated with the validated markers, their direct roles in blanchability remain unclear. Nevertheless, the gene *protein kinase superfamily*, through phosphorylation cascades, may regulate the expression of genes involved in cell wall biosynthesis and adhesion, but further functional characterization studies are required to elucidate these regulatory mechanisms.

Haplotype analysis across the promising loci, Ah05, Ah06 and Ah17 provides valuable insights for haplotype-based breeding. Specifically, the superior high-blanchability haplotypes (*Ah05HapBL3*, *Ah06HapBL5*, *Ah06HapBL10* and *Ah17HapBL6*) offer promising targets for introgression into breeding programs. Our findings conclude that integrating these superior high-blanchability haplotypes from the sub-species *fastigiata*, and agronomic type Spanish bunch or Valencia bunch into breeding programs will facilitate the development of groundnut cultivars with high-blanchability independent of agronomic type. A suggested cross between high-blanchability genotypes (ICG9507 and ICG297) might result in improved blanchability (80– 90%) **(Figure 5**). Moreover, ICG297 has also found to be a promising donor for blanchability along with other agronomic traits [9]. The required blanchability percentage, however varies depending on manufacturer specifications and end-product demands. For example, genotypes exhibiting a high percentage of split seeds are more suitable for products like candies and groundnut butter, where easy removal of the testa is advantageous. In contrast, low-blanchability genotypes are preferred for beer nuts and coated seed products, as the retained testa contributes to desired texture and facilitates better adherence of confectionery coatings. Thus, both high and low-blanchability traits are valuable depending on the processing application. This dual demand emphasizes the importance of aligning breeding objectives with specific end-use requirements, advocating for a more nuanced and market-driven approach to trait selection and cultivar development. This haplotype-based breeding strategy aligns with the goal of breeding groundnut varieties that meet the stringent quality standards required by the processing industry, thereby adding value to groundnut production chains.

## 5. Conclusion

Blanchability, the ease of seed coat removal post-roasting, is a critical trait for groundnut processing quality for a number of food applications. This study successfully integrated WGRS-based GWAS, and haplo-pheno analysis to identify the genetic architecture underlying blanchability. A total of 26 significant STAs were detected, with major loci on chromosomes Ah05, Ah06, and Ah17. Among these, the Ah17 chromosome explained, showing the highest phenotypic variance (PVE 54.03%). The STAs on Ah05, Ah06, and Ah17, emerged as a key genomic region, and revealed superior high-blanchability haplotypes *Ah05HapBL3, Ah06HapBL5/ Ah06HapBL10, and Ah17HapBL6*. These superior high-blanchability haplotypes predominantly originated from *fastigiata* subspecies; and agronomic types; Valencia bunch and Spanish bunch types; highlighting the influence of genetic background and ecogeographic adaptation. Conversely, low-blanchability haplotypes were mostly confined to *hypogaea* subspecies and agronomic types; Virginia runner and bunch types, indicating genetic constraints. Further, validation of two KASP markers, *SnpAH00567* and *SnpAH00569*, confirmed reproducibility across platforms, while challenges were encountered with the Ah17 validation point to the need for alternative marker strategies. Candidate genes near significant STAs, including *galactoside 2-alpha-L-fucosyltransferase-like protein, protein kinase superfamily* and *glycerophosphoryl diester phosphodiesterase*, suggest involvement in cell wall remodelling and seed coat adhesion, and hence can be targeted for future functional characterization. The targeted introgression of favourable haplotypes into low-blanchability germplasm offers a promising strategy for developing groundnut cultivars with improved processing quality independent of genetic background and agronomic type. Overall, this study provides a promising framework for haplotype-based breeding to customize blanchability and thereby enhancing the economic value of groundnut in global markets.

## Data availability statement

The phenotypic data used in this work is provided in Supplementary Table S1. The sequencing data generated in this study is deposited in NCBI with bio-project ID PRJNA1002116, PRJNA490835, PRJNA490832.

## Institutional Review Board Statement

Not Applicable

## Informed Consent Statement

Not Applicable

## CRediT authorship Contribution Statement

**Manish K. Pandey:** Conceived the idea, supervised and finalized the manuscript. **Kuldeep Singh and Ramachandran Senthil:** Contributed seed material and in seed multiplication of the mini-core collection. **Pasupuleti Janila:** Contributed the breeding lines for marker validation. **Priya Shah:** Phenotyped mini-core collection, performed the analysis and wrote the manuscript. **D. Khaja Mohinuddin, Ragavendran Abbai and Sunil S. Gangurde:** performed the analysis and reviewed, edited and improved the manuscript. **Madhvi Sharma, Prashant Singam**, **Chuanzhi Zhao, Sandip K. Bera, Mei Yuan, Xingjun Wang and Rajeev K. Varshney:** Reviewed, edited and improved the manuscript. All authors have read and agreed to the published version of the manuscript.

## Acknowledgments

Priya Shah acknowledges Joint Council of Scientific and Industrial Research-University Grant Commission (CSIR-UGC), Government of India for the award of fellowship for a Ph.D. and ICRISAT for providing research facilities.

## Supplementary Figures

Figure S1-S6.

## Supplementary Tables

Tables S1-S11.

