## Supplementary Figures S1-S6 for "Genomic analysis uncovers unique haplotype signatures from subspecies and agronomic types associated with blanchability in groundnut"

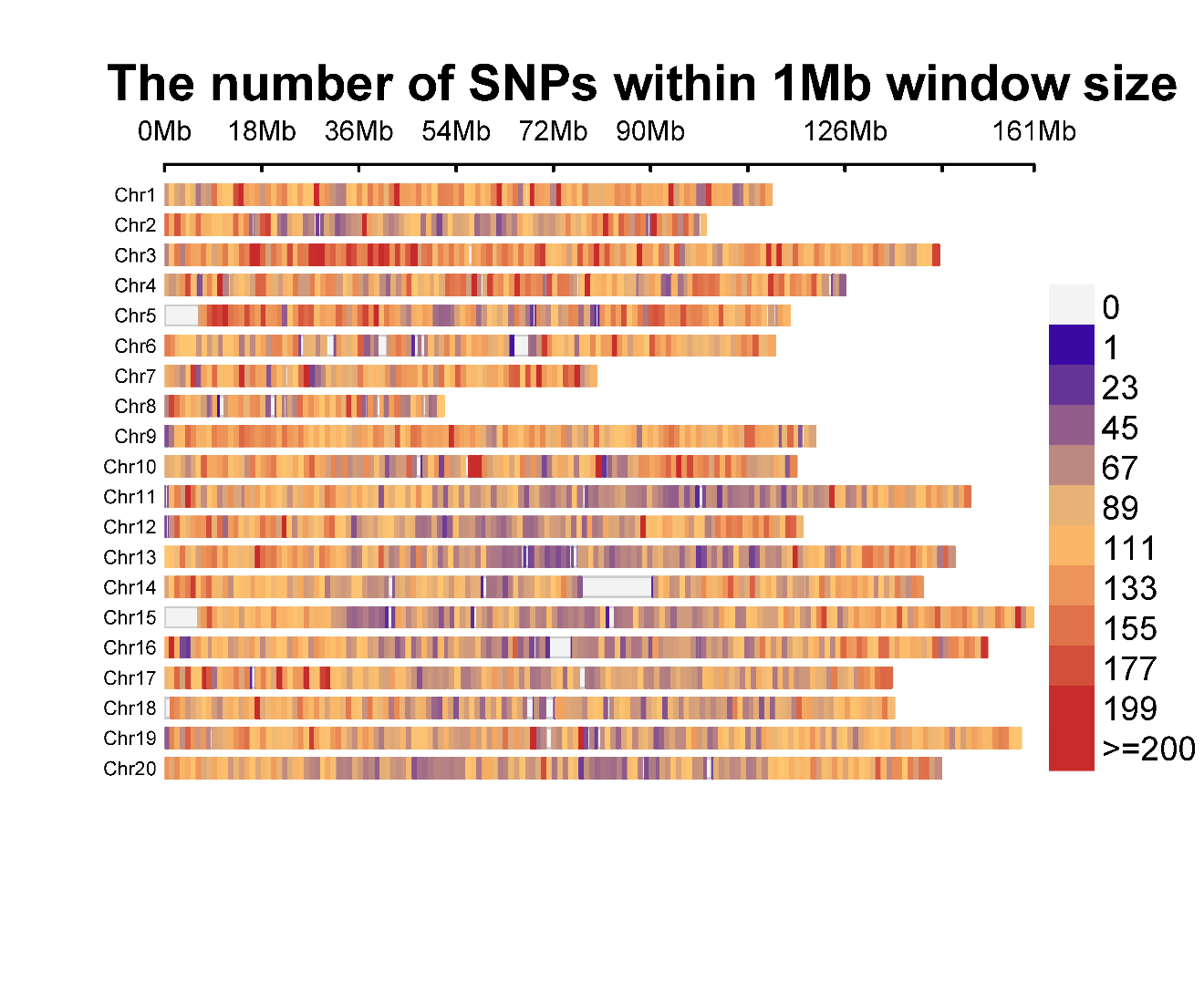

**Supplementary Figure S1: SNP density plot with 1Mb window size for WGRS data.** Red-colored regions, indicating the highest SNP density, appear sparsely, particularly on Chr Ah03 and Chr Ah05. The blue colored regions indicate low SNP density, whereas there are also white regions on chromosomes, signifying regions with low or no detected SNPs.

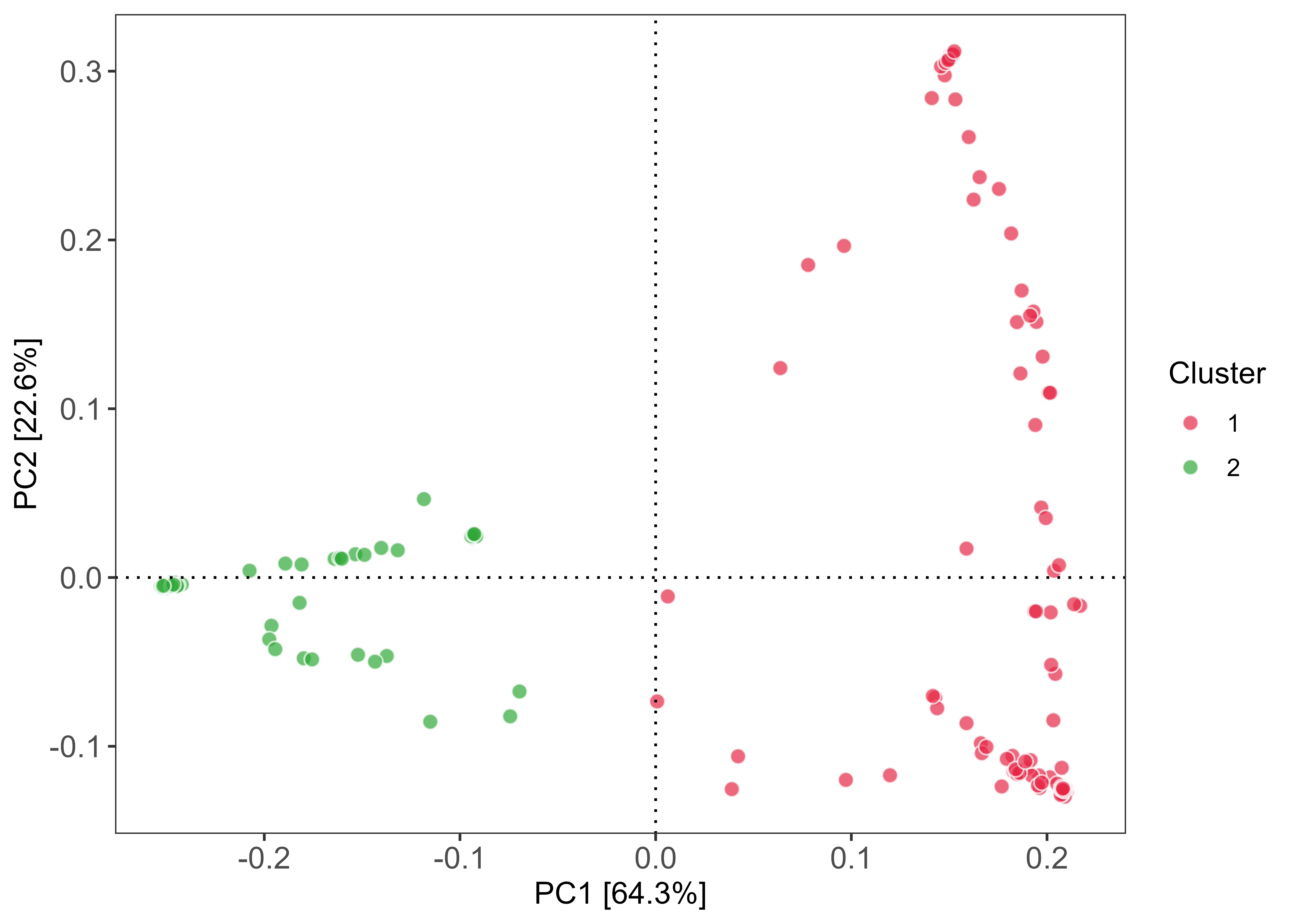

**Supplementary Figure S2: PCA for groundnut mini-core collection depicting (PC1+PC2) 86.9% phenotypic variability, and identified 2 distinct clusters (pink and green) based on groundnut sub-species, botanical variety and agronomic type.** Cluster1 (pink color): (88 genotypes): *fastigata*, valencia bunch/ hypogaea, virginia runner/ vulgaris, spanish bunch; Cluster2 (green color): (72 genotypes): *hypogaea*, virginia bunch or virginia runner/ hirsuta/ aequatoriana/ peruviana/ vulgaris, spanish bunch/ fastigata, valencia bunch.

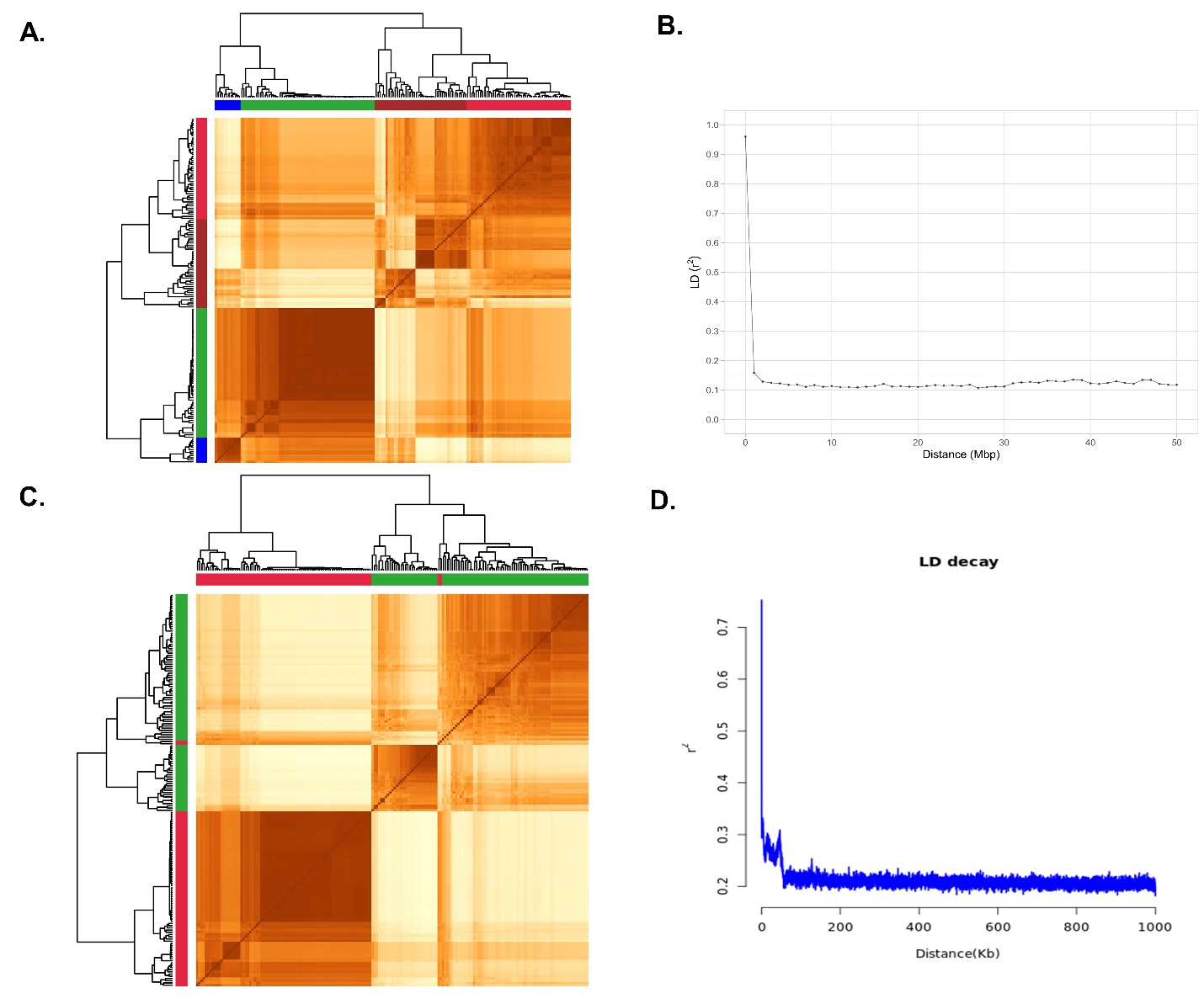

A.

B.

**Supplementary Figure S3: PCA, Population Structure analysis and genome-wide linkage disequilibrium,** A. Heat map of population structure, B. LD decay for groundnut minicore collection identified

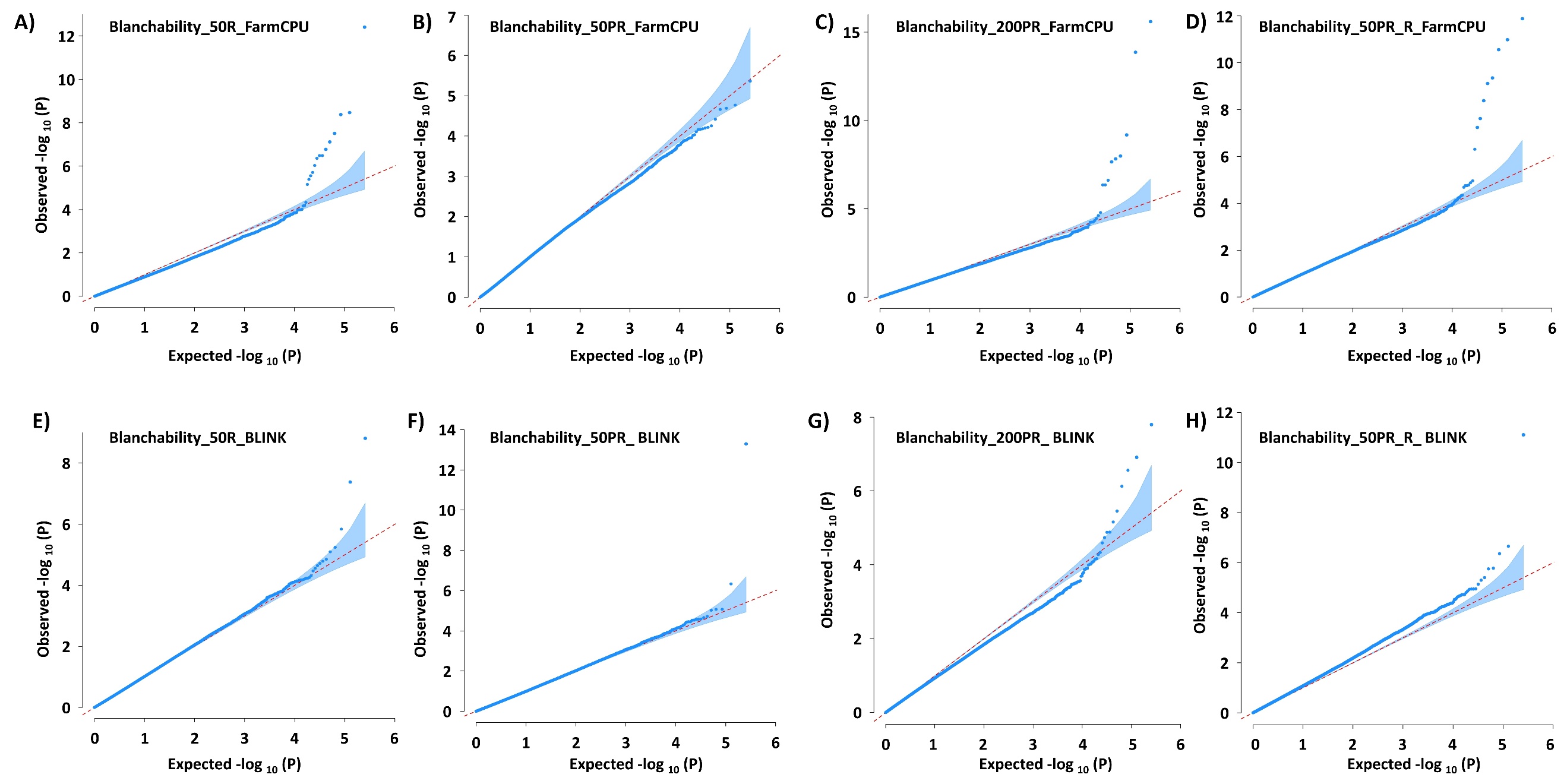

**Supplementary. Figure S4:** **Quantile-Quantile (Q-Q) plot representing the identified STAs associated with the blanchability on the basis** A, B, C, D: Q-Q plot representing the identified STAs associated with the blanchability on the basis FarmCPU and E, F, G, H: BLINK, model in GAPIT, using WGRS data for Blanchability 50R, 50PR, 200PR and 50PR_R, respectively.

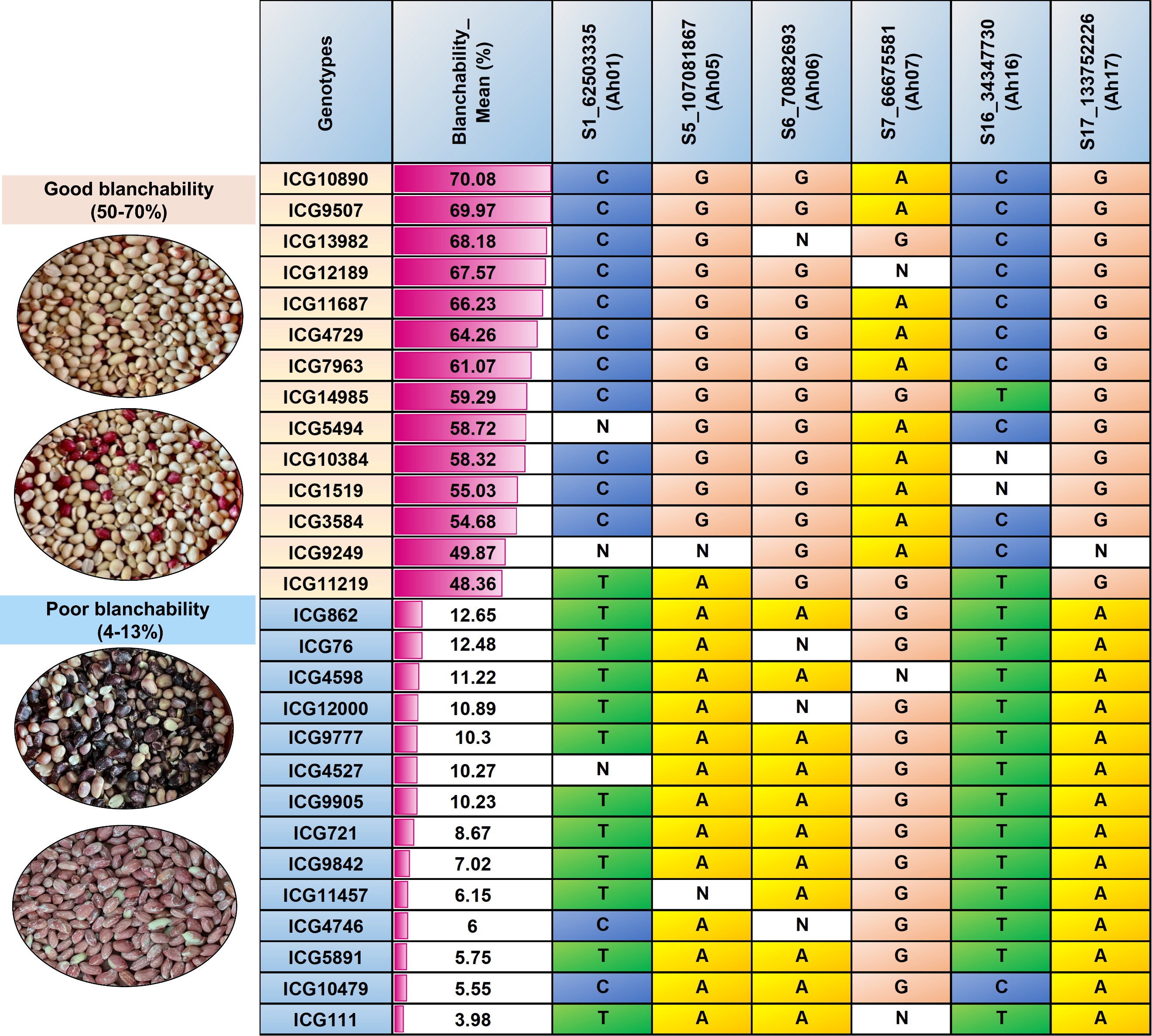

**Supplementary Figure S5: Six stable signiﬁcant polymorphic SNPs for blanchability in minicore collection,** including; S1_62503335, S5_107081867, S6_70882693, S7_66675581, S16_34347730, and S17_133752226.

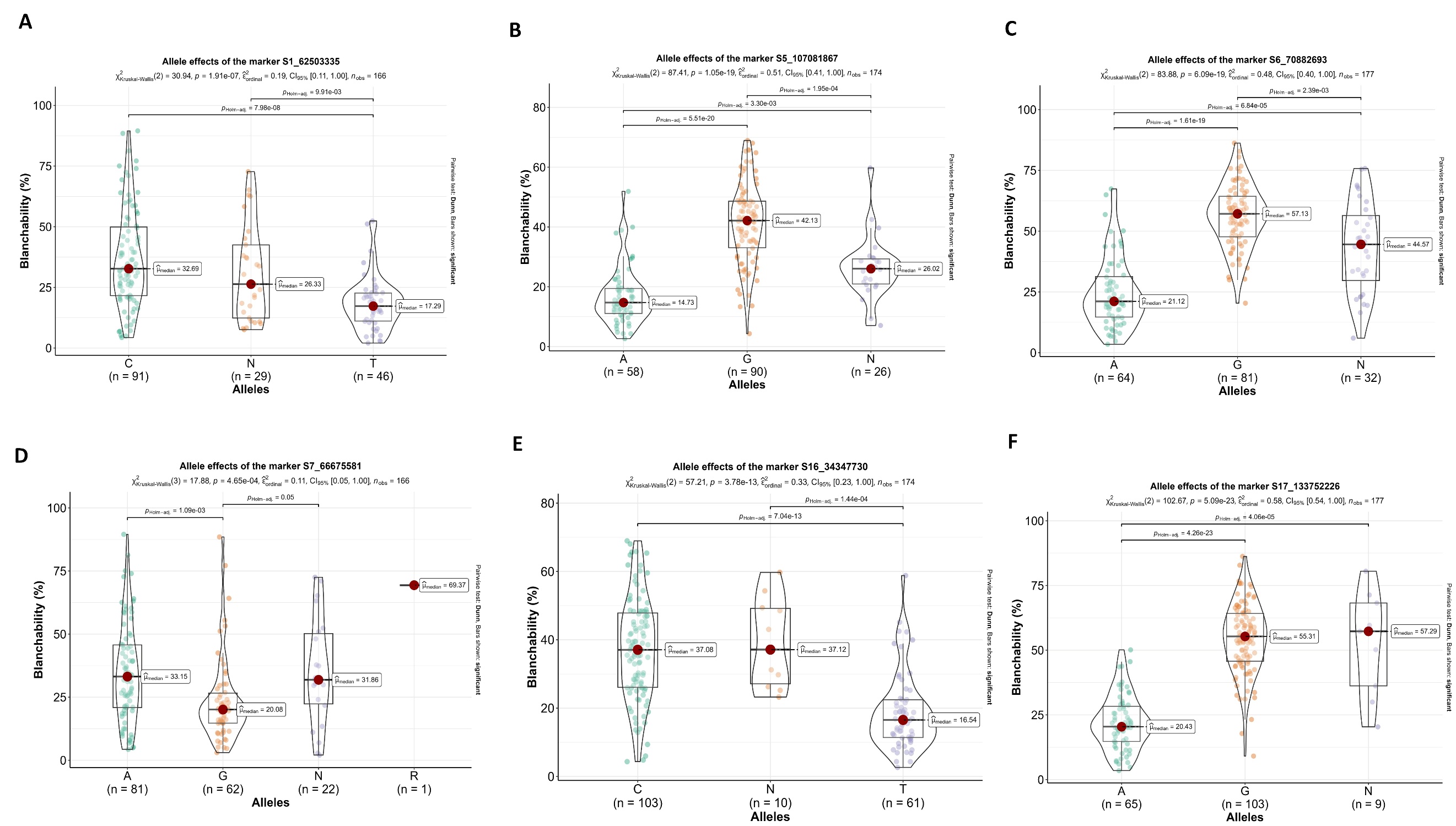

**Supplementary Figure S6: Allele-effect analysis for six stable signiﬁcant SNPs** including A. S1_62503335, B. S5_107081867, C. S6_70882693, D. S7_66675581, E. S16_34347730, F. S17_133752226. The plot depicts the number of the alleles for each of the six signiﬁcant SNPs in mini-core collection, and the contribution of these alleles to the phenotypic variation observed blanchability.
